# VCFdbR: A method for expressing biobank-scale Variant Call Format data in a SQLite database using R

**DOI:** 10.1101/2020.04.28.066894

**Authors:** Tanner Koomar, Jacob J Michaelson

## Abstract

As exome and whole-genome sequencing cohorts grow in size, the data they produce strains the limits of current tools and data structures. The Variant Call Format (VCF) was originally created as part of the 1,000 Genomes project. Flexible and concise enough to describe the genetic variations of thousands of samples in a single flat file, the VCF has become the standard for communicating the results of large-scale sequencing experiments. Because of its static and text-based structure, VCFs remain cumbersome to parse and filter in an interactive way, even with the aid of indexing. Iterating on previous concepts, we propose here a pipeline for converting VCFs to simple SQLite databases, which allow for rapid searching and filtering of genetic variants while minimizing memory overhead. Code can be found at https://github.com/tkoomar/VCFdbR

## Introduction

The Variant Call Format is a well-specified and flexible way to describe genetic variants identified by sequencing, along with annotations describing those variants. These annotations are a particularly powerful feature of the VCF, and set it apart from more efficient formats of expressing genetic variants, like PLINK and BGEN. Unfortunately, despite advances in processing VCFs [3] as a flat text file, searching and filtering variants in a VCF based on its annotations is slow and cumbersome.

When studying genetics of larger cohorts, it is often necessary to begin with exploratory analyses – which require rapid iteration and changes to filtering schemes. Data from these studies are often so large that they cannot be read directly into RAM, and writing subsets of variants to a hard disk repeatedly is slow and introduces overhead that does not scale well. Therefore, faster methods of identifying variants of interest from sequencing data are needed, such as databases or datastores.

For example, the GenomicsDB format, a datastore developed by GATK team and Intel [5], is fast, scaleable and efficient. However, it is currently only widely used for the merging of gVCFs produced by GATK, and much work is still needed to make it a standard that can replace the VCF. The Hail data-analysis library [10] does a good job of implementing GenomicsDB, but attempts to be an end-to-end solution. This requires learning its own interface and is not compatible with the tools many computational geneticists already use – particularly R [9] and Bioconductor [4] packages.

The Gemini database [8] is a somewhat more flexible alternative that takes the approach of converting a VCF to an SQLite database that can be manipulated with any suitable interface. SQLite is a single-file database that does not require a server, making it suitable for working on high performance computing clusters that are available and widely used by computational geneticists in academic institutions. However, Gemini compresses genotypes to save space, which slows down queries. Users can opt not to compress genotypes with Gemini, but this creates 1 column per sample (i.e. ‘wide’ format), which can rapidly exceed the column limit of an SQLite database. Additionally, this is not scaleable to very large cohorts which might result in databases which violate file system limits on the size of single file.

Many researchers would benefit from a method for converting a VCF to an SQLite database that is suitable for large cohorts. This requires genotypes to be stored in the more database-friendly ‘long’ format as well as an option to save genotypes to external files to avoid hitting single-file size limits. Such a method should also avoid operations that require writing intermediate files to disk and the requirement to learn bespoke interfaces. The R language is widely used by computational geneticists, and has the excellent general-purpose SQL database interface provided by the dbplyr package [11]. Additionally, R is under-served in this realm, as both Gemini and Hail utilize Python-based interfaces. For these reasons, we propose a relatively simple pipeline for converting a VCF to an SQLite database, which we call VCFdbR.

## The VCFdb Table Schema

### Overview

SQLite is a server-less, single-file relational database composed of one or more ‘tables’, which may be linked by shared ‘keys’. In a VCFdb created by VCFdbR, this ‘key’ is a integer that uniquely identifies each variant in the VCF. Specific columns within a given SQLite table may also be indexed, so that they may be searched rapidly.

Information describing each variant of a VCF is divided among up to three tables of a VCFdb. Core information and INFO columns are stored in a table called `variant_info`. If annotations produced by the Variant Effect Predictor (VEP) [6] are present, they are in a table called `variant_impact`. Genotypes are stored in a `variant_geno` table if the database is created in ‘Table-GT mode’, or as individual files in a separate directory if created in ‘File-GT mode’. Table-GT databases are simpler, producing a single portable database file. File-GT databases require care to move after creation, but allow for a database to be created for tens or hundreds of thousands of individuals (assuming sufficient disk space).

### Core Tables and Files

#### variant info

The `variant_info` table contains all of the core information that uniquely describes a variant. The columns that appear here depend on the annotations provided in the input VCF. To save disk space, it is recommended to remove all unneeded INFO columns from the VCF, e.g. with `bcftools annotate --remove`. Each variant appears in this table exactly once.

**Table 1.** variant_info table schema.

| variant_id | chr | start | end | ref | alt | qual | filter | an | ac | ... |
| --- | --- | --- | --- | --- | --- | --- | --- | --- | --- | --- |
| 1 | chr22 | 16050075 | 16050075 | A | G | 100 | PASS | 50 | 8 | ... |
| 2 | chr22 | 16050115 | 16050115 | G | A | 100 | PASS | 50 | 12 | ... |
| ... | ... | ... | ... | ... | ... | ... | ... | ... | ... | ... |
| 961 | chr22 | 16136747 | 16136751 | TATTA | T | 100 | PASS | 50 | 22 | ... |
| ... | ... | ... | ... | ... | ... | ... | ... | ... | ... | ... |

#### variant_impact

The `variant_impact` table is created if the input VCF has been annotated with VEP (i.e. it has an INFO field named CSQ). This information is placed into its own table because each variant may have multiple effects or affect multiple genes/transcripts. Thus, each variant may appear in `variant_impact` multiple times.

**Table 2.** variant_impact table schema.

| variant_id | consequence | impact | symbol | ... |
| --- | --- | --- | --- | --- |
| 840 | upstream_gene_variant | MODIFIER | LA16c-60H5.7 | ... |
| 840 | intron_variant | MODIFIER | NBEAP3 | ... |
| 840 | non_coding_transcript_variant | MODIFIER | NBEAP3 | ... |
| ... | ... | ... | ... | ... |

### Genotypes

#### Table-GT: variant_geno

When a VCFdbR is run in ‘table’ mode, the resulting Table-GT database will have a table called `variant_geno`. Each entry of this table describes the genotype (as dosage) of one individual for one variant, along with additional columns for all FORMAT fields from the VCF. With a moderate to large number of samples, this portion of the database takes up the most disk space. To save substantial space, it is recommended to remove all unneeded FORMAT fields from the VCF before database creation, e.g. with `bcftools annotate --remove`!. If the input VCF contains *n* samples and *m* variants, then this table will have rows equal to *n × m*.

**Table 3.**
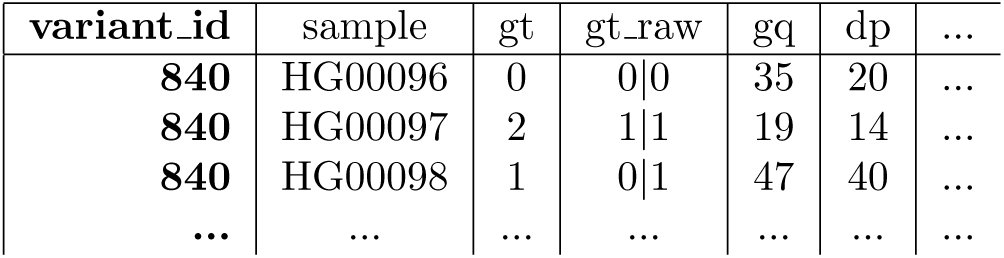
variant_geno table schema.

#### File-GT: geno directory

When a VCFdbR is run in ‘table’ mode, the resulting FILE-GT database will not have a `variant_geno`, instead one file per variant will be saved to a write-protected directory. The absolute path to these files will be saved to a `geno` column on the `variant_info` table. These files may be read in to R with the `read.rds()` or `read_rds()` commands, and otherwise have an identical layout to the genotypes of a Table-GT database. This separation of the genotypes from the annotations (which remain in the database) makes it possible to move the database to a faster but smaller disk for quicker filtering (e.g. a local SSD) while keeping the genotypes on a larger be slower disk (e.g. a network-connected hard disk).

**Table 4.** variant_info table schema for a File-GT database.

| variant_id | chr | start | ... | geno |
| --- | --- | --- | --- | --- |
| 1 | chr22 | 16050075 | ... | /path/prefix-genos/1.rds |
| 2 | chr22 | 16050115 | ... | /path/prefix-genos/2.rds |
| ... | ... | ... | ... | ... |
| 961 | chr22 | 16136747 | ... | /path/prefix-genos/961.rds |
| ... | ... | ... | ... | ... |

### Additional Table and Files

#### Miscellaneous Tables

Additional tables will be created based on the full content of the VCF header. For example, a sample table will list all samples, while an INFO table will contain the descriptions for each INFO column provided in the input VCF.

#### gene_map

If VEP annotations were provided on the input VCF, an additional `gene_map` table will be produced in the final database. This is a terse mapping of all of the genes and genomic elements which actually appear in the database. This table can be useful for identifying the universe of genes or other annotations which are present in the database.

**Table 5.** gene_map table schema.

| symbol | source | canonical | ... | feature_type |
| --- | --- | --- | --- | --- |
| LA16c-4G1.3 | Ensembl | YES | ... | Transcript |
| NBEAP3 | Ensembl | YES | ... | Transcript |
| ... | ... | ... | ... | ... |

#### Genomic Ranges

After database creation, VCFdbR creates a GenomicRanges representation of each variant. The GenomicRanges data structure is the bedrock of many Bioconductor genomics packages. This file can be used to identify variants of interest based on annotations external to the database, or even to generate new columns to be added to the `variant_info` table. This is a minimal representation of the variants, and only includes the position, REF/ALT alleles, and possibly the path to the corresponding variant files (for File-GT databases only).

## Database Creation

### Input Requirements

Input VCFs should conform to the format standard, be gzipped/tabixed indexed, and must have multi-allelic sites split, e.g. with

~~~
bcftools norm -m -
~~~

A VCF which has been annotated with VEP should be passed through

~~~
sed /^#/\! s/;;/;/g
~~~

to ensure proper parsing of the annotations when read into R. Finally, VCFdbR requires an up-to-date version of R, preferably greater than version 3.5. Specific package requirements can be found on the GitHub repo.

### Pipeline Overview

To accommodate RAM limits, VCFdbR operates by reading in chunks of the input VCF and writing to the database iteratively. To identify chunks of variants, minimal information from the VCF is read into R with the VariantAnnotation [7] package, and GenomicRanges corresponding to 1,000 variants each are written to a file. The number of variants per chunk can be reduced based on memory constraints at the expense of some processing time.

The header of the VCF is then parsed and written to the miscellaneous tables of the database. For each chunk, the VCF’s INFO field is then parsed and searched for a CSQ field. Then, the genotypes are parsed, converting them to ‘long’ format. Once the data INFO, CSQ and FORMAT are fully parsed, they are appended to the corresponding database tables (`variant_info, variant_impact` and `variant_geno`, respectively). If operating in File-GT mode, the parsed FORMAT fields are written to the file system instead of the database.

After all VCF chunks have been processed, the database tables are indexed. Indexing always includes the `variant_id`, and new indexes can be added by the user at any time. Finally, the GenomicRanges representation of the database is written to disk and the `gene_map` table is created (if the input VCF had VEP annotations).

## Benchmarks

### Database Creation

For benchmarking, three databases were prepared from phase 3 of the 1,000 Genomes Project [1]. First, a File-GT database from the whole genome sequences of all 2,504 subjects from the project was created. Then, both File-GT and Table-GT databases were created from the exome-captured variants for first 1,000 samples. A summary of the time required for each of the major steps of database creation (chunking, building, and indexing) can be seen in Table **??**. All databases were created with a chunk size of 10,000, and the File-GT databases were written with four threads. This – and all other benchmark steps – were carried out on server-grade compute nodes and NFS-networked disks on the high performance computing cluster at the University of Iowa.

| Dataset | Samples | Variants | VCF Size | Mode | Componant | Size | Step | Time |  |
| --- | --- | --- | --- | --- | --- | --- | --- | --- | --- |
| exome | 1,000 | 1,417,043 | 693 MB | table |  |  | chunk |  | 2 min |
|  |  |  |  |  | database | 22.6 GB | build | 1 hr | 32 min |
|  |  |  |  |  | geno files |  | index |  | 20 min |
|  |  |  |  |  | **total** | **22.6 GB** | **total** | **1 hr** | **54 min** |
| exome | 1,000 | 1,417,043 | 693 MB | file |  |  | chunk |  | 2 min |
|  |  |  |  |  | database | 2.8 GB | build | 2 hr | 3 min |
|  |  |  |  |  | gene files | 34.4 GB | index |  | 12 min |
|  |  |  |  |  | **total** | **37.2 GB** | **total** | **2 hr** | **17 min** |
| genome | 2,504 | 85,211,481 | 29.7 GB | file |  |  | chunk | 2 hr | 19 min |
|  |  |  |  |  | database | 64.8 GB | build | 218 hr | 8 min |
|  |  |  |  |  | geno files | 2.55 TB | index | 5 hr | 56 min |
|  |  |  |  |  | **total** | **2.61 TB** | **total** | **225 hr** | **23 min** |

### Variant Filtering

Because the filtering of variants based on annotations of interest occurs entirely within the database, it can be extremely fast. This requires all annotations used for filtering to be indexed, but is generally worth the time and small increase in database size since the time to query an indexed column is on the log scale.

To illustrate this, the time to filter variants to those with an alternate allele frequency less than 5% and within each gene in both the genome and exome databases prepared from 1,000 Genomes Phase 3 is plotted in Figure 1. Although this requires filtering the variants on both the `variant_impact` (for gene) and `variant_info` (for alternate allele frequency) tables, genes with 10,000 qualifying variants typically take less than 1.5 seconds to identify and read into memory (Figure 1.B).

**Figure 1.**
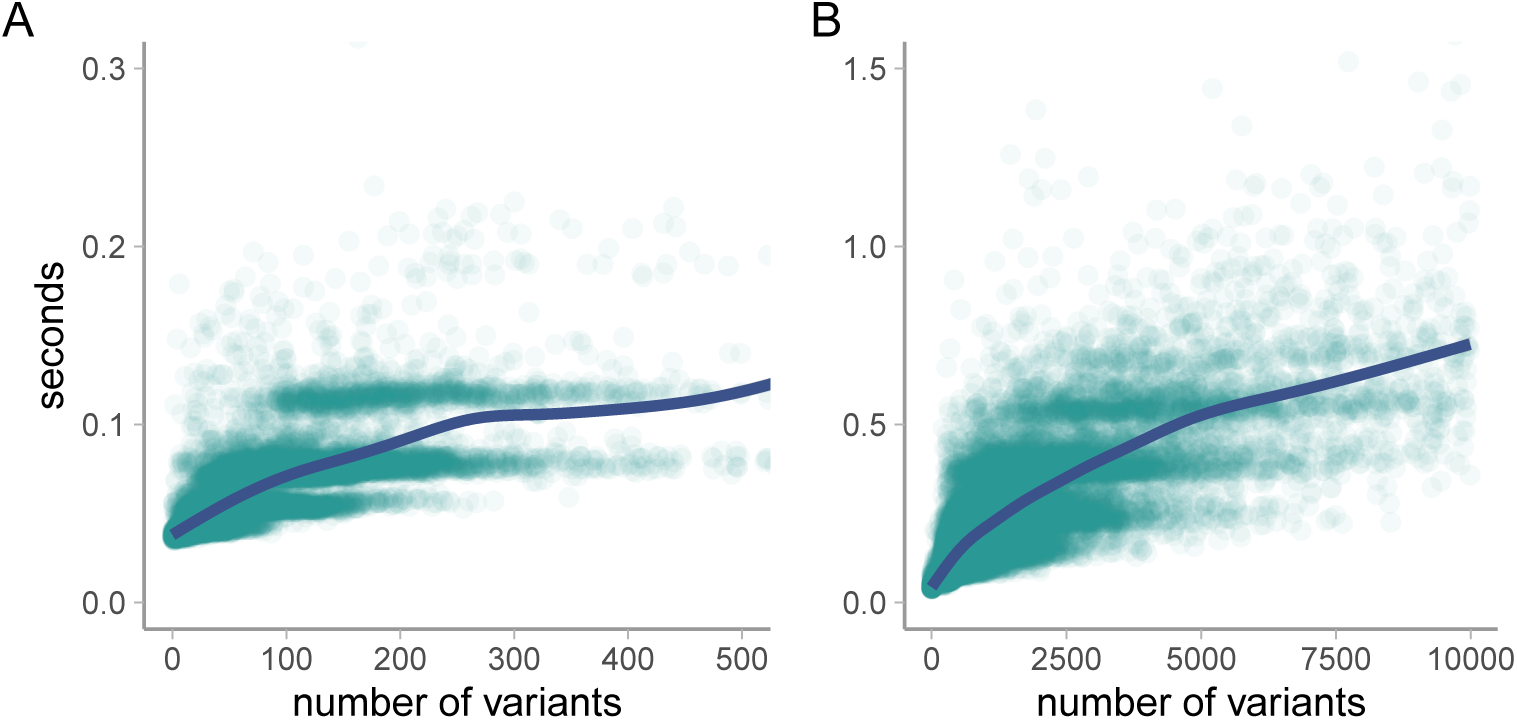
The time to identify variants of interest based on indexed columns is nonlinear with the number matching variants. The filtering here is performed on both numeric (allele frequency) and text (gene name) columns in both exome data (**A**) and whole-genome (**B**) data.

Because this example performs no filtering based on genotypes or other FORMAT fields, there is no effective difference between File-GT and Table-GT databases. If substantial filtering is needed on FORMAT fields, there is a significant increase in the speed of filtering for Table-GT databases (provided the `variant_geno` table is appropriately indexed).

### Genotype Pulling

Though computationally trivial, reading genotypes into memory is by far the most I/O intensive part of working with a VCFdb. This process can be sped up somewhat through the use of multiprocessing across more than one processor core. However, disk and network limitations mean that using more threads may have diminishing returns even before the principles of Amdhal’s Law [2] become relevant.

To illustrate this, random subsets of variants from each of the three 1,000 Genomes databases described above were generated, ranging in size from 50 to 5,000 variants. The genotypes for these variant sets were then read in to memory using a range of cores for multiprocessing. The time taken for each subset of variant across each database and number of multiprocessing cores is illustrated in Figure 2. On the particular hardware used for these benchmarks, there was little improvement in speed when using more than 4 threads to read genotypes simultaneously, and virtually no difference between using 8 and 16 threads. This indicates that disk or network speeds likely saturated when using around 4 cores. Performing this sort of benchmarking is recommended when running on new hardware to ensure resources are used as efficiently as possible.

**Figure 2.**
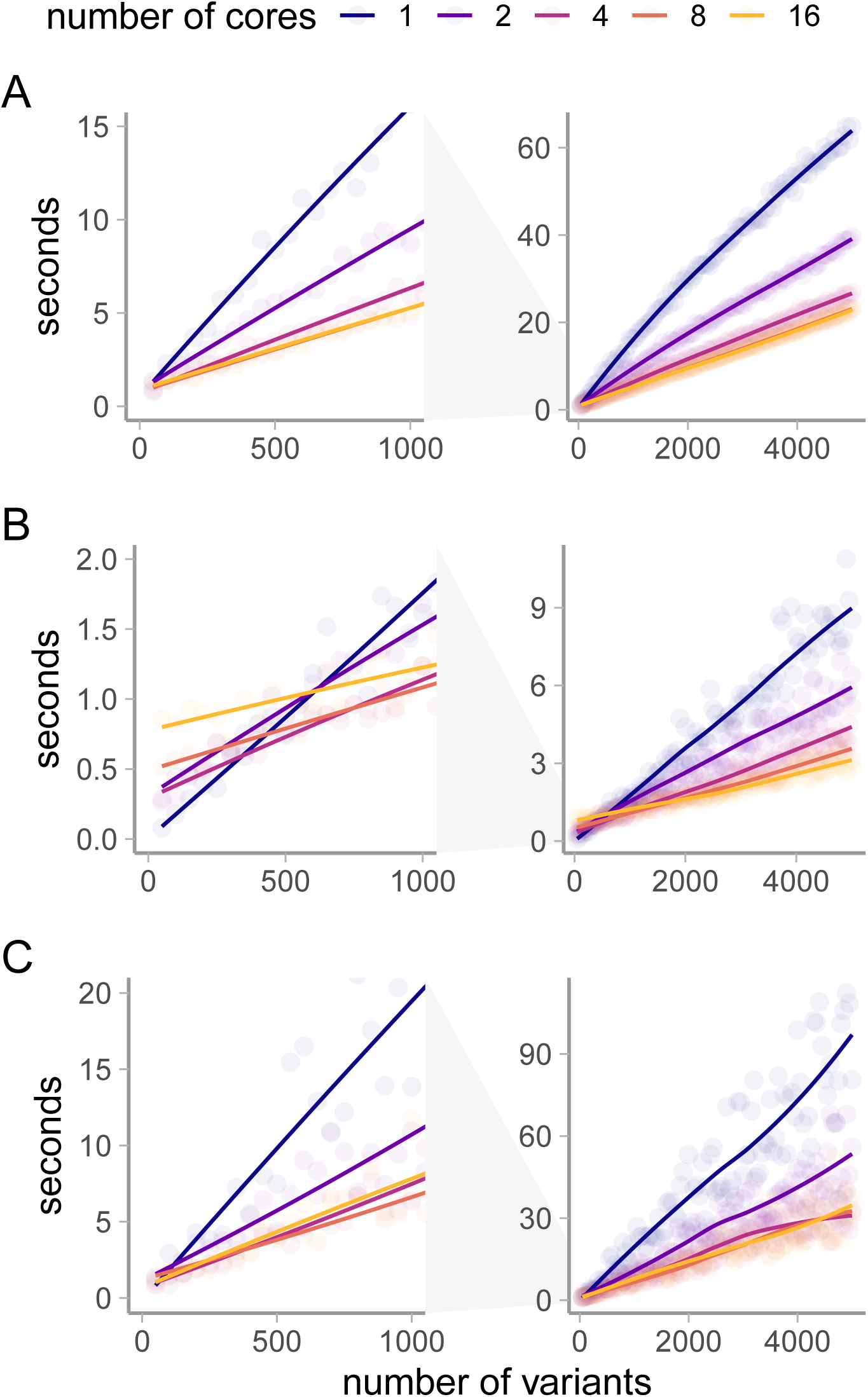
Reading genotypes with many simultaneous processes has limited benefit. In benchmarks of (**A**): a Table-GT exome database with 1,000 samples, (**B**): a File-GT exome database with 1,000 samples, and (**C**): a File-GT whole genome database with 2,504 samples, speed gains from using more than 4 simultaneous processes is minimal.

## Acknowledgments

Ethan Bahl, Leo Brueggeman, Yongchao Huang, Taylor R. Thomas, and everyone else who tested and used VCFdb through multiple iterations.

